# Hybrid Clustering of Long and Short-read for Improved Metagenome Assembly

**DOI:** 10.1101/2021.01.25.428115

**Authors:** Yakang Lu, Lizhen Shi, Marc W. Van Goethem, Volkan Sevim, Michael Mascagni, Li Deng, Zhong Wang

## Abstract

Next-generation sequencing has enabled metagenomics, the study of the genomes of microorganisms sampled directly from the environment without cultivation. We previously developed a proof-of-concept, scalable metagenome clustering algorithm based on Apache Spark to cluster sequence reads according to their species of origin. To overcome its under-clustering problem on short-read sequences, in this study we developed a new, two-step Label Propagation Algorithm (LPA) that first forms clusters of long reads and then recruits short reads to these clusters. Compared to alternative label propagation strategies, this hybrid clustering algorithm (hybrid-LPA) yields significantly larger read clusters without compromising cluster purity. We show that adding an extra clustering step before assembly leads to improved metagenome assemblies, predicting more complete genomes or gene clusters from a synthetic metagenome dataset and a real-world metagenome dataset, respectively. These results suggest that hybrid-LPA is a good alternative to current metagenome assembly practice by providing benefits in both scalability and accuracy on large metagenome datasets.

**Availability and implementation:** https://bitbucket.org/zhong_wang/hybridlpa/src/master/.

## 1 INTRODUCTION

Metagenomics offers a fast track to directly study the microbial communities in their natural habitat without laboratory cultivation (Tyson et al., 2004; Hugenholtz and Tyson, 2008). Next-generation DNA sequencing (NGS) technologies have greatly expedited metagenomic discoveries, yielding deep insights into the composition, structure, and dynamics of complex microbial communities (Arumugam et al., 2011; Hess et al., 2011; Xu, 2006). Driven by the rapid development of NGS experimental technologies and modern, scalable metagenome assemblers, large numbers of individual microbial genomes can now be readily assembled from a single experiment or from meta-analyses constituting large cohorts of metagenomic datasets (Stewart et al., 2019; Parks et al., 2017; Nayfach et al., 2020). Currently, the Illumina Sequencing Platform is the predominant NGS platform for metagenome sequencing due to its high-throughput, low-cost, and high accuracy (average error rate <1%), despite that its short read length creates limitations on some downstream analysis tasks such as gene discovery (Wommack et al., 2008), read classification, or genome assembly (Breitwieser et al., 2019). To overcome these limitations, various strategies have been developed to either create synthetic long reads by assembly (Zimin et al., 2013) or experimentally (such as Moleculo, White et al. (2016)), but these methods bring additional experimental and/or computational costs.

Single-molecule, long-read sequencing technologies developed by Pacific Biosciences (PacBio, Eid et al. (2009)) and Oxford Nanopore Technologies (ONT, Schneider and Dekker (2012)) have been successfully applied to single-genome sequencing projects, yielding very high-quality genome assemblies from microbes to human (Chin et al., 2013; Koren and Phillippy, 2015; Logsdon et al., 2020; Sevim et al., 2019). These long reads, up to 100kb in length, can effectively resolve large repeats or structural variations that pose challenges to short-read based assemblers. Long-read sequencing has not been widely adopted in metagenome sequencing, however, mainly because of two reasons. Firstly, PacBio and ONT long reads have error rates as high as 30% (Eid et al., 2009; Schneider and Dekker, 2012). These errors, predominantly small insertions and deletions (indels), make the assembly process difficult and error-prone if they are not corrected. Secondly, compared with short-read sequencing, these technologies, when applied to complex metagenome projects, incur higher costs and lower throughput.

Recently, hybrid approaches have emerged to take advantage of the complementary characteristics of short and long-read sequencing technologies. Combining the high accuracy of short-read sequencing and the high read length of long-read sequencing, some genome assemblers such as Unicycler (Wick et al., 2017) and hybridSPAdes (Dmitry et al., 2016) showed promising results for single-genome assembly. However, most popular metagenome assemblers, including MEGAHIT (Li et al., 2015), MetaSPAdes (Nurk et al., 2017) and MetaHipmer (Hofmeyr et al., 2020), do not support hybrid assembly yet. The feasibility and potential benefits of a hybrid strategy in metagenome assembly were recently demonstrated by leveraging long reads for a second-round assembly of contigs from those metagenome assemblers (Bertrand et al., 2019).

We previously developed a scalable metagenome clustering tool called SpaRC (Shi et al., 2018; Li et al., 2020) based on Apache Spark. SpaRC can form pure and complete clusters with long-read sequencing technologies. However, it tends to produce a large number of small clusters on short-read datasets (under-clustering) unless multiple samples from the same community are available. To illustrate this point, Table 1 shows the results of running SpaRC on two short-read datasets, each derived from a single sample of a synthetic microbial community: BMock12 (Sevim et al., 2019) and CAMI2 Simulated Toy Human Gut Metagenome (Sczyrba et al., 2017; Bremges and McHardy, 2018). In both experiments, SpaRC generated pure clusters but their completeness was very low.

**Table 1.** Clustering Performance on Single-sample, Short-read, Synthetic Metagenome Datasets.

|  | # reads | # clusters | Median Purity | Median Completeness |
| --- | --- | --- | --- | --- |
| BMock12 | 10,517,108 | 79,915 | 100 | 0.29 |
| CAMI2 Simulated<br>Toy Human | 296,027,232 | 1,347,826 | 100 | 11.62 |

Motivated by the success of the above mentioned hybrid assemblers, in this study we explored a hybrid approach for metagenome read clustering to overcome the under-clustering problem of SpaRC. As SpaRC’s core algorithm is based on the Label Propagation Algorithm (LPA), we first experimented three alternative label propagation strategies after long reads were added. Next, we explored the effect of using different proportions of long reads since long-read sequencing is relatively more costly. We also compared hybrid clustering performance of long-read datasets from both PacBio and ONT platforms. Finally, we evaluated the impact of hybrid clustering on downstream genome assembly and gene-cluster discovery performance, using a synthetic and a real-world metagenome dataset, respectively.

## 2 MATERIALS AND METHODS

### 2.1 The Hybrid-LPA algorithm

SpaRC uses Label Propagation Algorithm (LPA) originally proposed by Raghavan (Raghavan et al., 2007) to partition the read graph (Shi et al., 2018). Briefly, the algorithm begins by initializing each read with a unique label, followed by iteratively updating the label of each node to the label of the majority of its neighbors. After several iterations or until no further label propagation is possible, densely connected groups of reads are partitioned into clusters. LPA is capable of resolving genomes with shared reads and has near linear computational performance. SpaRC can be run at two different modes: “local mode” only cluster reads based on their overlap, while “global mode” further clusters the results from local mode based on multiple sample statistics (Li et al., 2020).

Here we explored three strategies for hybrid clustering with both long- and short-reads (Figure 1A):

**Figure 1.**
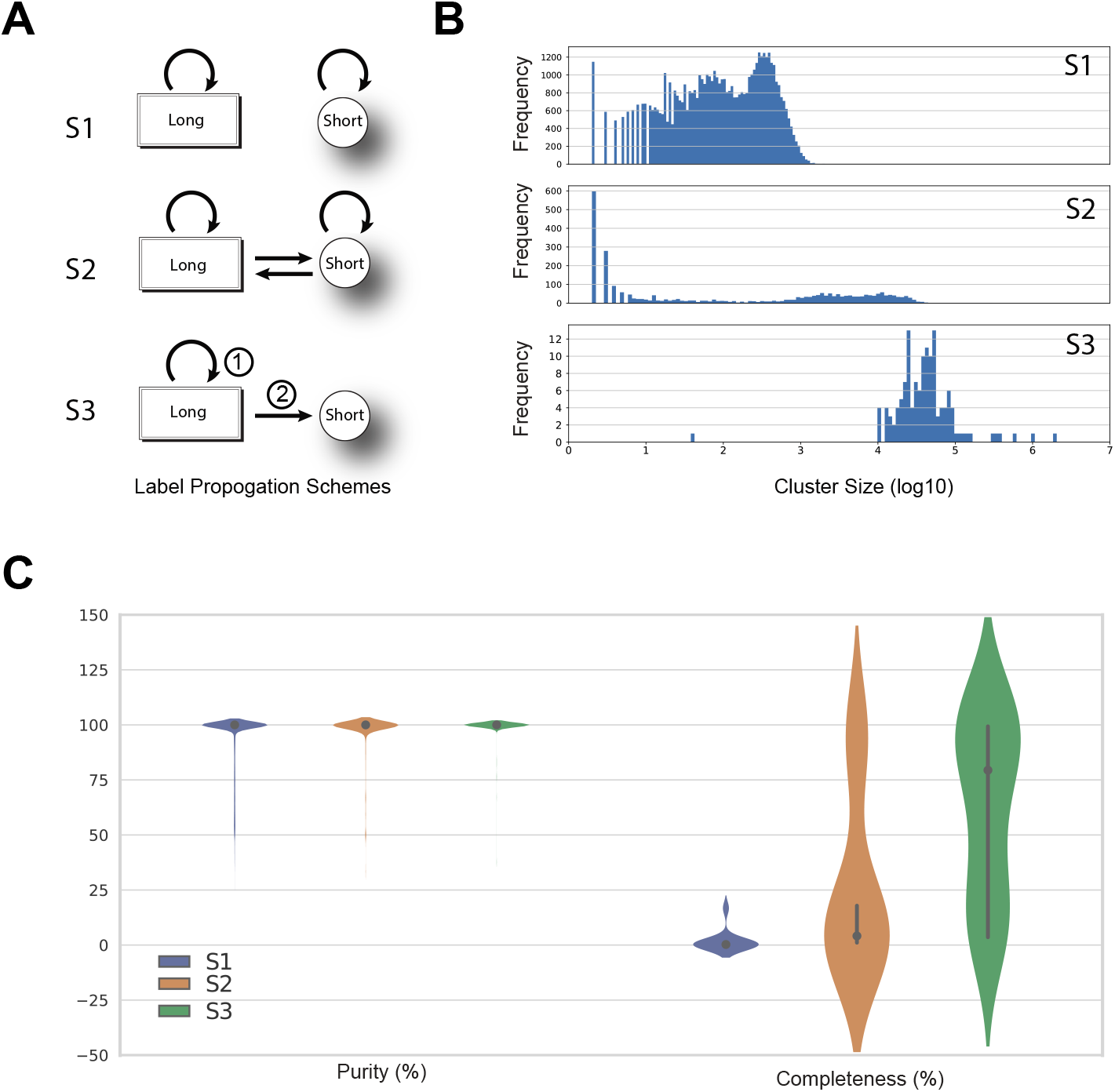
**(A)** Three alternative clustering strategies for hybrid-LPA. (S1) “Additive” strategy: clustering labels can only propagate among long reads or among short reads, respectively. No propagation was allowed between long and short reads. (S2) “Mixed” strategy: labels can be propagated among both long and short reads indiscriminately. (S3) “Long-then-short” strategy: in the first step, labels were only allowed to propagate among long reads, then they were propagated to short reads. No propagation was allowed among short reads. **(B)** A comparison of three label propagation strategies on cluster size improvement on the BMock12 dataset. The number of clusters (*Y-axis*) at each cluster size in log10 (*X-axis*), from top to bottom: S1, S2, S3. **(C)** A comparison of three label propagation strategies on the purity and completeness of clusters on the BMock12 dataset. Violin plots of purity and completeness distributions are shown in percentage (*Y-axis*).

- In the first “additive” strategy (S1), cluster labels can only propagate among long reads or among short reads, respectively. No propagation is allowed between long and short reads. This was done by running SpaRC at local clustering mode separately on the short-read and long-read datasets, and then combine the clustering results.
- In the second “mixed” strategy (S2), labels are allowed to propagate among both long and short reads indiscriminately: labels can propagate from long to long, short to short, long to short or *vice versa*. This was done by first combining the short- and long-read datasets, followed by running SpaRC at local clustering mode.
- In the third “long-then-short” strategy (S3), initially labels are only allowed to propagate among long reads. After all long reads finish updating their labels, their labels are allowed to propagate to short reads. This new algorithm, hereafter referred as hybrid-LPA, was implemented in both MPI and UPC++ in order to fit different HPC environments.

### 2.2 Datasets and Data Preprocessing

The BMock12 (Sevim et al., 2019) dataset was derived from a mock community that consists of 12 bacterial strains with genome sizes ranging 3.2 to 7.2 Mbp. One of the bacterial species in the set, *M. coxensis*, has a negligible number of reads in the dataset, therefore, BMock12 effectively contains 11 bacterial strains. The reads from BMock12 were downloaded from the NCBI Sequence Read Archive (SRA) using accessions SRX4901586 (ONT), SRX4901584 (PacBio set 1), SRX4901585 (PacBio set 2), and SRX4901583 (Illumina). Table 2 lists the statistics of these datasets. The Illumina short-read dataset from this community was pair-end sequenced at 150bp. The two ends were concatenated by an “N” (resulting a 301bp fragment) before being fed into SpaRC. In this paper, we took 5% of the reads from the original dataset to conduct the experiments.

**Table 2.** Sequencing data statistics.

| Dataset | Statistics | Illumina | ONT | PacBio |
| --- | --- | --- | --- | --- |
| BMock12 | #Reads | 211,448,444 | 187,507 | 389,806 |
|  | #Bases | 63,384,840,109 | 3,737,495,058 | 2,583,337,248 |
|  | Max Length | 301 | 145,720 | 45,165 |
|  | Min Length | 301 | 120 | 50 |
|  | Avg Length | 301 | 19,932.6 | 6,627.2 |
|  | Median Length | 301 | 17,900 | 5,800 |
|  | Std.Dev | / | 11,225.3 | 4,283.2 |
| Biocrust | #Reads | 141,172,036 | / | 20,042,887 |
|  | #Bases | 37,368,694,112 | / | 111,977,437,956 |
|  | Max Length | 301 | / | 138,853 |
|  | Min Length | 35 | / | 50 |
|  | Avg Length | 264.7 | / | 5586.9 |
|  | Median Length | 301 | / | 5427 |
|  | Std.Dev | 51.5 | / | 3,324.0 |

The Biocrust dataset was derived from a biological soil crust sample collected from Moab, UT, USA. Biocrusts are specialized microbial communities consisting of primary producers, such as cyanobacteria, mosses, and lichens, and associated heterotrophs. They are aggregated organosedimentary communities that colonize and stabilize the soil surfaces of arid environments, preventing soil erosion and promoting nutrient status by fixing both atmospheric carbon and nitrogen (Van Goethem et al., 2021). The two ends of a Illumina short-read pair are 151 and 150bp. The two ends were merged by BBMerge (Bushnell et al., 2017). The merged Illumina reads, as well as the PacBio reads, were masked for low-complexity sequences by BBDuk using default parameters (sourceforge.net/projects/bbmap/). The resulting fragments were used as input for SpaRC.

### 2.3 Running SpaRC and Hybrid-LPA

Small-scale experiments in this work were performed on the Amazon Web Services (AWS) Cloud. Apache Spark (ver 2.3.1) services and Hadoop (ver 2.8.4) are provided by the Elastic MapReduce (EMR) on AWS. Specifically, we first used SpaRC to generate read graphs (EMR, emr-5.17.0). Then we used one node (r4.16xlarge) with 64 CPU cores and 488GB memory to run hybrid-LPA. On the EMR cluster, one node is used as the master and all other nodes (r4.2xlarge) are used as workers. Depending on the size of the input datasets, various number of workers are used (20 workers for BMock12 and 200 for the Biocrust dataset).

Large-scale experiments were performed on Berkeley Lab’s High-performance Computing system (Lawrencium, https://sites.google.com/a/lbl.gov/hpc/) and Department of Energy’s National Energy Research Scientific Computing Center (NERSC, https://www.nersc.gov/). In these environments, SpaRC jobs were run on standalone Spark clusters created on-demand. Specifically, 600 Cori KNL nodes (each has 68 physical cores and 96 GB of memory) on NERSC were used for the Biocrust dataset.

### 2.4 Metagenome Assembly and Binning, Biosynthetic Gene Cluster Prediction

For the BMock12 dataset, each cluster from the hybrid-LPA output was assembled by metaSPAdes (ver 3.13.1) using default parameters (Nurk et al., 2017). Contigs from all clusters were combined for binning with MetaBAT 2 (Kang et al., 2019) using default parameters. In the assembly-only method, raw reads were assembled with metaSPAdes followed by binning with MetaBAT 2. MetaQuast (version 5.0.2) was used to evaluate assembly quality for both two methods (Mikheenko et al., 2016).

From the assembled biocrust metagenomes (performed using metaSPAdes, Canu (Koren et al., 2017) and metaFlye (Kolmogorov et al., 2020), providing 3 assemblies) we deduplicated the contigs using BB-Dedup using default parameters(sourceforge.net/projects/bbmap/) to only include unique sequences by removing redundant contigs. All contigs longer than 5 kb were retained for secondary metabolite production using antiSMASH v5.2.0 under strict settings to preclude the detection of false-positives (Blin et al., 2019). Here, biosynthetic gene clusters (BGCs) were retained if they were longer than 5 kb after manual inspection of the domain architecture. Finally, we compared the quantity of unique BGCs detected when clustering-then-assembling to assembly-only (metaFlye assembly only, as it produced the largest number of BGCs).

## 3 RESULTS

### 3.1 Long reads increase clustering performance

To test whether or not combining long reads with short reads improved clustering performance, we designed three strategies (Materials and Methods) to include long reads in SpaRC’s LPA step (Figure 1A). We ran the three strategies on the synthetic BMock12 dataset (Materials and Methods) with 12 known genomes and used three metrics to measure read clustering performance: read cluster size (number of reads in a cluster), purity (percent of reads from the predominant genome in a cluster) and completeness (percent of reads from the predominant genome in a cluster). For these experiments, we used Illumina short reads and ONT long reads. Since we aimed at exploring how the long reads help with short reads clustering, these metrics were calculated based on short reads only. In addition, as we did not expect SpaRC to distinguish different strains of the same species, strain-level differences were ignored when clustering purity was calculated.

Figure 1B illustrates the cluster size comparison between these different label propagation strategies. The additive strategy (S1) produces many small clusters. Clusters formed from the mixed strategy (S2) showed a bi-modal size distribution, characterized by the presence of many larger clusters and small clusters. In contrast, the long-then-short strategy (S3) only produces a small number of clusters, most of them are very large. These strategies resulted in similar numbers of short reads in clusters (Table 3). However, the number of clusters was reduced from 85,398 (S1) to 136 (S3), while the mean cluster size was increased from 125.3 (S1) to 75,749.1 (S3). Consequently, the median completeness was increased from 0.25% (S1) to 79.42% (S3). As shown in Figure 1C, this increase of genome completeness by S3 was reflected in that the majority of clusters having better completeness, a significant shift from the other two strategies. These improvements in cluster size and completeness did not come with a decreased clustering purity, with a median purity 100% and a mean purity 99.65% (Figure 1C). Clustering performance of the long-then-short strategy also outperforms the mixed strategy in terms of completeness, number of clusters, and cluster size.

**Table 3.** Clustering performance comparison between the three LPA strategies.

|  | #clusters | #reads | % of reads clustered | mean cluster size | median completeness | median purity | mean purity |
| --- | --- | --- | --- | --- | --- | --- | --- |
| S1 | 85,398 | 10,312,376 | 96.38 | 125.3 | 0.25 | 100 | 97.90 |
| S2 | 15,145 | 10,312,376 | 96.38 | 706.5 | 4.13 | 100 | 94.62 |
| S3 | 136 | 10,298,222 | 96.25 | 75,749.1 | 79.42 | 100 | 99.65 |

These results suggest that long reads can greatly improve metagenome read clustering performance and that the hybrid clustering strategy presented here is an effective way to solve the under-clustering problem with metagenomic short reads.

Although long reads greatly reduce the under-clustering problem in the above experiment, they did not solve the over-clustering problem, as some clusters contain reads from different genomes. Among the top 20 largest clusters, 17 of them are pure clusters at the species level (Table 4). The biggest cluster, consisting 2 million reads (20% of the clustered reads), mixed reads from two different closely-related species (*Marinobacter sp*.*1* and *Marinobacter sp*.*8*) of the same genus (*Marinobacter*), owing to the fact that these species have an average nucleotide identity (ANI) of 78.1%, and they share 105,617 common 31-mers, making them difficult to be distinguished (Supplemental Table S1). As expected, the clustering algorithm could not distinguish closely related strains of the same species, such as *Halomonas sp. HL-4* and *Halomonas sp. HL-93*, with 3,126,579 shared 31-mers and an ANI of 98.5%. This pair of genomes spread 14 of the top 20 clusters. Different species with a large number of shared k-mers could also get clustered together, as the second and third largest clusters each contain multiple genomes. Some of these genomes are related, but some are not clearly indicating an over-clustering problem.

**Table 4.**
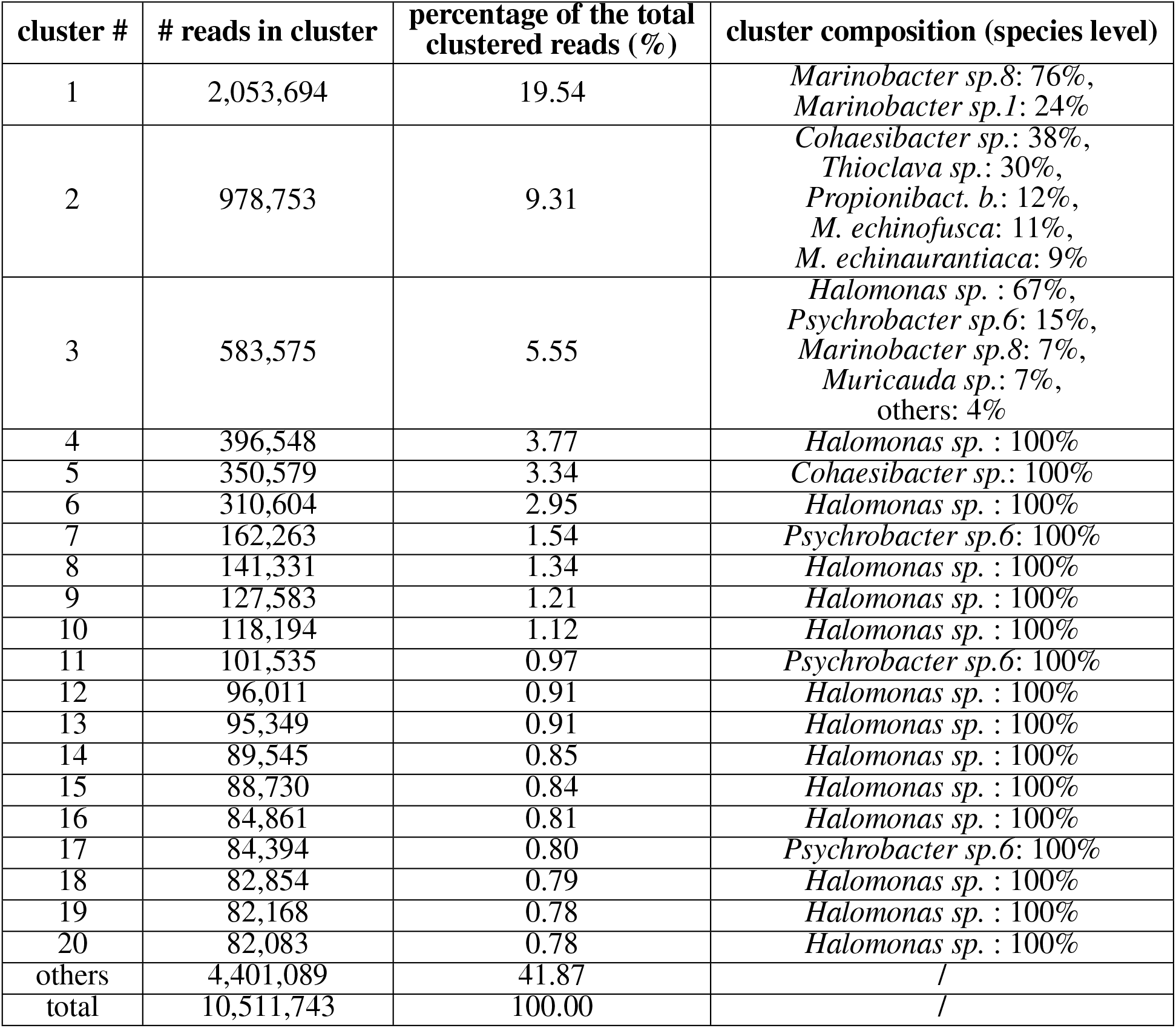
Top 20 cluster size and composition.

### 3.2 Small amounts of long-read data sufficiently improve clustering

As long-read sequencing technologies have higher cost and lower throughput, we tested whether or not limited numbers of long reads can help short-read metagenome clustering. In the following experiments done on the BMock12 dataset, we gradually increased the amount of ONT reads added to Illumina reads and compared the hybrid clustering performance.

As shown in Figure 2, adding just 1% ONT reads already produces a pronounced effect, increasing the mean cluster size to over 50,000 reads. Except for some variations when below 10% of the ONT reads were added, adding more ONT reads increases the mean cluster size, even though the increase gets smaller. The total number of clusters first rises, then steadily falls after 5% ONT reads. The total number of clustered short reads remains largely unchanged. As we added more long reads (>10% of total), the number of reads clustered, the number of clusters formed, and the mean cluster size all become stable. These results suggest a small fraction of long reads can significantly improve short read clustering, and the hybrid clustering approach could be a cost-effective metagenome clustering method.

**Figure 2.**
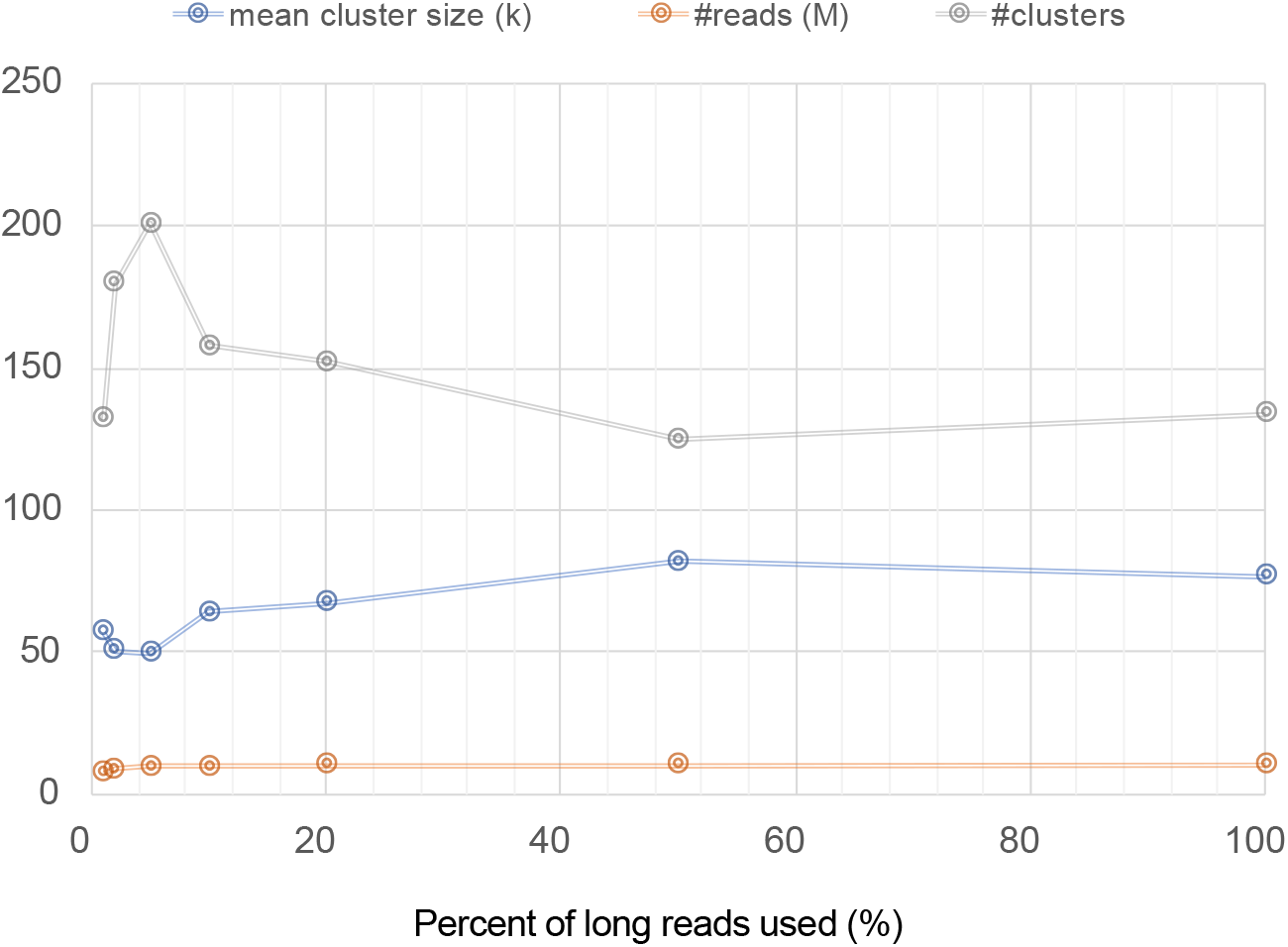
The effect of different amounts of ONT reads added to Illumina short reads on cluster size: the number of clusters (blue line), the number of reads being clustered in millions (M, grey line), and the mean cluster size in thousands (K, orange line) vary as different percentages of ONT long reads are added (*X-axis*)

### 3.3 Read length, not the sequencing platform, has a major impact on the cluster size

In theory, longer read lengths should increase the clustering performance, as their ability to bridge short reads gets better with length. To test this hypothesis, we added shorter PacBio reads from the same BMock12 dataset and compared the results to the above obtained from ONT reads. The read length distribution of ONT and PacBio reads is shown in Figure 3A.

**Figure 3.**
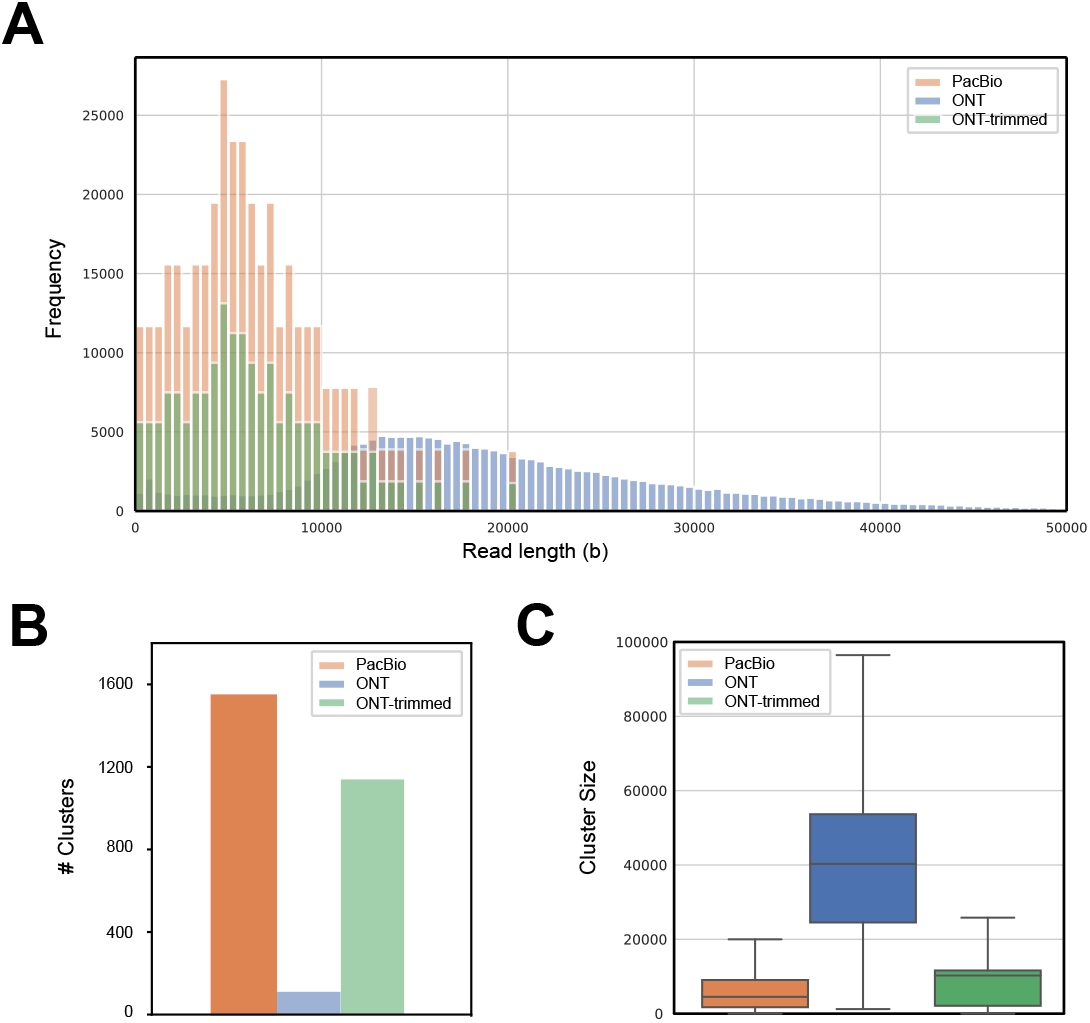
The dependency of hybrid clustering performance on read length. **(A)** Read length distribution of PacBio (orange), ONT (blue) and trimmed ONT (green) reads to match PacBio read length distribution in BMock12: the read length (*X-axis*) is plotted with its respective number of reads (*Y-axis*) for ONT and PacBio sequencing platforms. **(B)** The number of clusters from hybrid-LPA using PacBio, before and after trimming ONT. The number of clusters is much smaller in ONT than PacBio but becomes comparable after the trimming. **(C)** Box plots of cluster size from hybrid-LPA using PacBio, before and after trimming ONT. The cluster size (*Y-axis*) after trimming ONT read length is comparable to PacBio, both are much smaller than ONT

As expected, ONT read hybrid clustering gave much better results than those from PacBio reads. The number of clusters from the ONT experiment is 136, while the PacBio produced 1,502 clusters (Figure 3B). The corresponding genome completeness metrics were measured at 79.42% and 7.09% for ONT and PacBio, respectively. The size of the clusters produced by adding ONT reads is much larger than that of PacBio reads (Figure 3C). To investigate whether this difference is caused by different platforms rather than by different read lengths, we trimmed the ONT reads so that they have the same length distribution as the PacBio reads (Figure 3A) and then repeated the experiment. The number of clusters became 1,149 by adding the trimmed ONT reads, which is very similar to the results obtained from the PacBio reads. And the genome completeness for trimmed ONT was reduced to 10.91%. The cluster size distribution is also comparable to the results of PacBio experiment. In all three experiments, the median purity metrics of the clusters are comparable, ranging from 97.73%-100%. These results confirmed that the read length, rather than the long-read sequencing platform, has a major impact on clustering performance.

### 3.4 Hybrid clustering improves downstream metagenome assembly and gene cluster discovery

To investigate whether or not the improved clustering results produced by hybrid clustering can translate into better downstream applications, we used two common scenarios as examples. First, on the BMock12 dataset where the set of genomes are known, we asked whether or not hybrid clustering produces better metagenome-assembled-genomes (MAGs). Second, we used a real-world Biocrust metagenome dataset without known references (Materials and Methods), and asked whether or not hybrid clustering could produce more predicted biosynthetic gene clusters (BGCs), locally clustered genes that together encode a biosynthetic pathway for the production of secondary metabolites (Medema et al., 2015). In both cases we wanted to compare the results to metagenome assembly with hybrid clustering (hereafter we refer as “SpaRC-hybrid”) and without (“Assembly”). The steps in this two methods are otherwise identical except in the SpaRC-hybrid the assembly was done on the clusters instead of on raw reads. A schematic view of the two methods is shown in Figure 4A.

**Figure 4.**
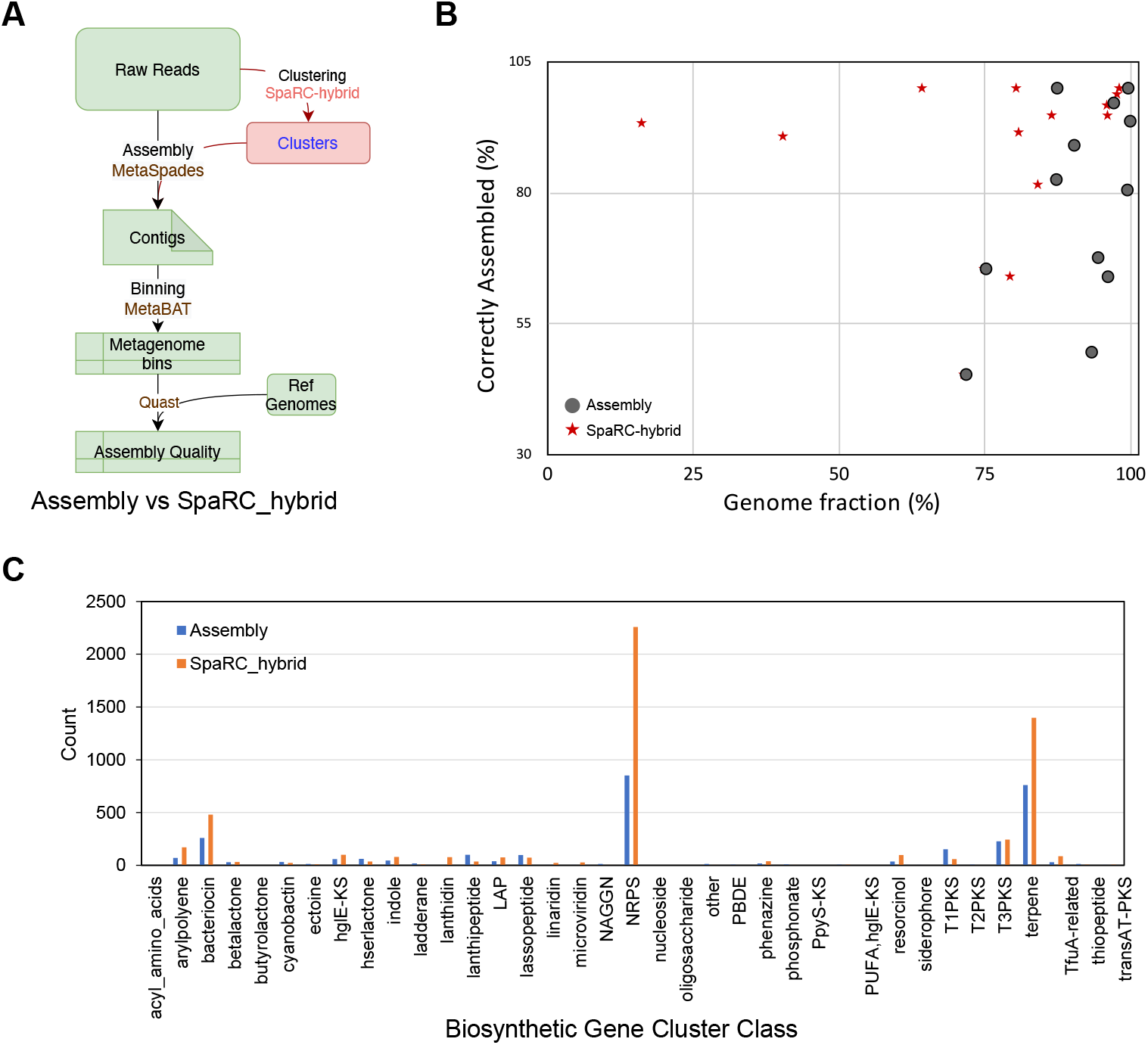
**(A)** A schematic view of metagenome hybrid assembly methods. Default “Assembly” method first assembles the raw reads (short and long) using an assembler (such as MetaSpades Nurk et al. (2017)), and then bins the resulting contigs into metagenome bins by a binner (such as MetaBAT Kang et al. (2015)). If reference genomes are available, the quality of the bins can be evaluated by Quast. The “SpaRC-hybrid” method first clusters the raw reads into clusters, then assembles the clusters into contigs, followed by the same procedures as the Assembly approach. **(B)** A comparison of assembled genome quality between the Assembly and SpaRC-hybrid approach on the BMock12 dataset. Two metrics measured by Quast, Genome Fraction percentage (*X-axis*) and percent of correctly assembled (*Y-axis*), are shown for each genome. Metrics for the Assembly method are shown in circles and the SpaRC-hybrid method in stars. **(C)** Bar charts of biosynthetic gene clusters (BGCs) predicted from the Biocrust dataset. Here we directly compared the difference in predicted BGCs counts for major BGC classes between assembly with metaFlye and our SpaRC-hybrid approach with the same assembler.

For the BMock12 dataset, the quality of genome bins were evaluated using Quast (Gurevich et al., 2013). Quast produces many metrics, here we focused on two assembly-related ones: the percent of genome coverage that measures the extent that a genome bin covers a reference genome, and percent of correctly assembled that measures the percent of assemblies aligned to references without any mis-assemblies (Figure 4B). Using 80% genome fractions and 90% correctness as cut-offs, the SpaRC-hybrid method produces 8 good genomes while the Assembly method only produced 4, supporting the notion that hybrid clustering improves downstream genome assembly. The full Quast report is available in Supplemental Table S2. Other differences between these two methods we noticed include SpaRC-hybrid producing much smaller N50s, higher rates of mismatches and small indels. These observations suggest the under-clustering problem still exists to some extent, so that the assemblers do not have sufficient read coverage for correcting the errors in long reads, or producing good contiguity.

For the Biocrust dataset, we used the ability to discover unique Biosynthetic Gene Clusters (BGCs) as a metric to test the benefit of hybrid-LPA over the Assembly method without prior clustering (Materials and Methods). Overall, the SpaRC-hybrid method predicted more BGCs than the Assembly method alone (Figure 4C). MetaFlye assembly derived from SpaRC-hybrid clusters gave 5,458 unique BGCs, considerably more than those from the Assembly approach (2,988 BGCs). In almost every category SpaRC-hybrid predicted more BGCs, with the most pronounced difference in Non-ribosomal peptides, a common and important class of secondary metabolites encoded by multidomain non-ribosomal peptide synthetases (NRPS). A complete list of the counts are available in Supplemental Table S3. The hybrid approach also predicted more complete gene clusters (i.e., it is not truncated on either of the contig edges) than the assembly-only approach, 1,100 vs 712 (Van Goethem et al., 2021). The longest NRPS is novel (based on sequence similarity to the entire NCBI nr database) and is a full-length gene cluster of 79,925 bp.

We made similar observations when we assembled the clusters using CANU instead of MetaFlye (Supplemental Table S3), suggesting hybrid clustering by SpaRC-LPA can benefit downstream assemblers in general.

## 4 DISCUSSION

In this work, we developed a new scalable algorithm, SpaRC-hybrid to incorporate long reads into metagenome read clustering. We showed that the hybrid clustering method can reduce the under-clustering problem in clustering experiments with only short reads. We also demonstrated that the read length, rather than the sequencing technologies, has a big impact on the clustering performance. Furthermore, improved clustering results can greatly augment downstream metagenome assembly or gene cluster discovery.

While SpaRC-hybrid can effectively leverage long reads to reduce the under-clustering problem in short reads, it does not reduce over-clustering problems, where similar genomes, or genomes sharing large genetic elements (horizontally transferred genes, very closely-related homologs, mobile elements, etc.) are clustered together. Given that SpaRC-hybrid uses long reads to build the initial read graph, it should alleviate the problem to some extent at this stage. However, over-clustering can still happen at the short-read recruitment stage. Using stringent read overlapping criteria may reduce the problem, but this may come with a cost of under-clustering and loss in sensitivity. In complex real-world metagenome datasets, this is unlikely to be a major drawback, as the overall complexity within a cluster could be greatly reduced compared to the original dataset. We may not be able to completely deconvolute a large, complex metagenome into single genomes, but can effectively partition into many simpler metagenomes. With the decreasing cost and increasing throughput of long-read sequencing, ultimately we may have to use only long reads for metagenome clustering to overcome the over-clustering problem.

Currently, SpaRC-hyrbid tends to produce a more fragmented assembly containing more small errors (mismatches, small indels). The most likely cause for this problem is under-clustering, as reads from the same genome were separated into different clusters. In the subsequent assembly step, each cluster does not have sufficient read coverage for good contiguity, precluding the building of contigs. Some additional matrices may be needed to further reduce under-clustering. The small errors are likely those carried over from long read sequencing. In the control dataset, BMock12, there are only 12 species, applying an error-correction step by either using short reads to correct long reads, or using long reads to correct each other, should improve this problem. In real-world complex metagenome datasets error-correction may not be reliable, especially those with a large strain-level diversity. The recent PacBio high-fidelity reads may be used to avoid small errors, but at the expense of read-length reduction and more under-clustering.

## CONFLICT OF INTEREST STATEMENT

The authors declare that the research was conducted in the absence of any commercial or financial relationships that could be construed as a potential conflict of interest.

## AUTHOR CONTRIBUTIONS

The study was conceived by ZW. LS implemented the hybridLPA algorithm. YL, LS, and VS performed the BMock data analyses. MWVG and LS performed the Biocrust analysis. All authors contributed to the writing and editing of the manuscript and approved the submitted version.

## FUNDING

The work was supported by the National Natural Science Foundation of China (No. 61802246) and the 111 Project (No. D18003). Marc W. Van Goethem, Volkan Sevim and Zhong Wang’s work was supported by the U.S. Department of Energy, Office of Science, Office of Biological and Environmental Research under Contract No. DE-AC02-05CH11231

## DISCLAIMER

No approval or endorsement of any commercial product by the National Institute of Standards and Technology is intended or implied. Certain commercial software, products, and systems are identified in this report to facilitate better understanding. Such identification does not imply recommendations or endorsement by NIST, nor does it imply that the software and products identified are necessarily the best available for the purpose.

## ACKNOWLEDGMENTS

We thank the scientific computing group at Berkeley lab, especially Dr. Shawfeng “Shaw” Dong and Dr. Wei Feinstein for their support to run SpaRC on Lawrencium. We thank members of NERSC, for their support to run SpaRC on the Cori system.

## Notes

### Competing Interest Statement

The authors have declared no competing interest.

