## Supplemental Table 1 for "Hybrid Clustering of Long and Short-read for Improved Metagenome Assembly"

### Table S1. Number of shared 31-mers and ANI between species

| species name | metrics | Muricauda sp. | Thioclava sp. | Cohaesibacter sp. | Propionibact. b. | Marinobacter sp.8 | Marinobacter sp.1 | Psychrobacter sp.6 | M. echinaurantiaca | M. echinofusca | M. coxensis | Halomonas sp. HL-4 | Halomonas sp. HL-93 |
|---|---|---|---|---|---|---|---|---|---|---|---|---|---|
| Muricauda sp. | # shared 31-mers | - | 117 | 137 | 57 | 101 | 101 | 82 | 58 | 58 | 58 | 109 | 109 |
| | ANI | - | 74.6% | 69.5% | 73.2% | 70.1% | 70.8% | 69.2% | 73.9% | 73.8% | 73.8% | 71.5% | 70.4% |
| Thioclava sp. | # shared 31-mers | - | - | 1,036 | 156 | 209 | 231 | 216 | 154 | 205 | 151 | 242 | 245 |
| | ANI | - | - | 71.8% | 70.6% | 70.0% | 70.0% | 71.3% | 70.5% | 70.1% | 70.4% | 70.1% | 70.1% |
| Cohaesibacter sp. | # shared 31-mers | - | - | - | 118 | 283 | 279 | 177 | 140 | 124 | 130 | 220 | 220 |
| | ANI | - | - | - | 69.0% | 70.5% | 69.9% | 71.9% | 69.6% | 69.2% | 69.8% | 69.8% | 70.0% |
| Propionibact. b. | # shared 31-mers | - | - | - | - | 64 | 71 | 96 | 3,686 | 3,135 | 3,359 | 71 | 71 |
| | ANI | - | - | - | - | 69.6% | 68.7% | 71.6% | 72.5% | 72.6% | 72.6% | 70.3% | 69.8% |
| Marinobacter sp. 8 | # shared 31-mers | - | - | - | - | - | **105,617** | 1,902 | 103 | 103 | 100 | 1,197 | 1,234 |
| | ANI | - | - | - | - | - | **78.1%** | 70.8% | 69.4% | 69.8% | 69.6% | 71.4% | 71.5% |
| Marinobacter sp. 1 | # shared 31-mers | - | - | - | - | - | - | 1,743 | 103 | 107 | 103 | 1,238 | 1,243 |
| | ANI | - | - | - | - | - | - | 71.1% | 69.3% | 69.3% | 69.5% | 71.5% | 71.5% |
| Psychrobacter sp.6 | # shared 31-mers | - | - | - | - | - | - | - | 102 | 102 | 102 | 453 | 445 |
| | ANI | - | - | - | - | - | - | - | 75.5% | 74.7% | 75.4% | 70.3% | 70.2% |
| M. echinaurantiaca | # shared 31-mers | - | - | - | - | - | - | - | - | **293,016** | **254,879** | 73 | 73 |
| | ANI | - | - | - | - | - | - | - | - | **85.7%** | **84.1%** | 70.7% | 70.1% |
| M. echinofusca | # shared 31-mers | - | - | - | - | - | - | - | - | - | **244,517** | 82 | 86 |
| | ANI | - | - | - | - | - | - | - | - | - | **83.8%** | 70.9% | 70.8% |
| M. coxensis | # shared 31-mers | - | - | - | - | - | - | - | - | - | - | 86 | 90 |
| | ANI | - | - | - | - | - | - | - | - | - | - | 69.9% | 69.8% |
| Halomonas sp. HL-4 | # shared 31-mers | - | - | - | - | - | - | - | - | - | - | - | **3,126,579** |
| | ANI | - | - | - | - | - | - | - | - | - | - | - | **98.5%** |
| Halomonas sp. HL-93 | # shared 31-mers | - | - | - | - | - | - | - | - | - | - | - | - |
| | ANI | - | - | - | - | - | - | - | - | - | - | - | - |
