## Supplemental Table 2 for "Hybrid Clustering of Long and Short-read for Improved Metagenome Assembly"

| Table S2: Comparison of assembly quality of BMock12 dataset | | | | | | | | |
|---|---|---|---|---|---|---|---|---|
| Method | genomeId | Genome fraction (%) | Correct (%) | Misassembled contigs length | # mismatches per 100 kbp | # indels per 100 kbp | N50 (bp) | Fraction>80% and Misassembly<10% |
| AssemblyOnly | 2615840527 | 99.56 | 100.00 | 0 | 1.79 | 0.22 | 2,783,390 | 1 |
| | 2615840533A | 93.21 | 49.59 | 481,197 | 9.74 | 2.2 | 481,197 | 0 |
| | 2615840533B | 96.04 | 63.99 | 1,361,938 | 1.61 | 0.48 | 770,690 | 0 |
| | 2615840601 | 97.06 | 97.17 | 140,307 | 3.35 | 0.16 | 271,614 | 1 |
| | 2615840646 | 99.92 | 93.69 | 283,374 | 13.36 | 11.24 | 534,540 | 1 |
| | 2615840697 | 87.21 | 100.00 | 0 | 5.24 | 0.82 | 2,219,840 | 1 |
| | 2616644829 | 94.33 | 67.65 | 1,354,434 | 46.79 | 1.81 | 659,299 | 0 |
| | 2617270709 | 99.42 | 80.55 | 620,253 | 10.31 | 2.81 | 2,346,159 | 0 |
| | 2623620557 | 87.06 | 82.55 | 1,246,442 | 2277.27 | 658.93 | 13,816 | 0 |
| | 2623620567 | 90.18 | 89.09 | 774,014 | 1749.03 | 406.09 | 13,816 | 0 |
| | 2623620609 | 0.00 | 0.00 | * | * | * | * | 0 |
| | 2623620617 | 71.37 | 45.33 | 1,641,088 | 1631.02 | 46.68 | 184,948 | 0 |
| | 2623620618 | 74.83 | 65.51 | 1,069,503 | 493.03 | 25.25 | 184,948 | 0 |
| SpaRC-hybrid | 2615840527 | 99.58 | 100.00 | 0 | 15.73 | 41.97 | 66,210 | 1 |
| | 2615840533A | 40.36 | 90.81 | 38,309 | 47.65 | 77.64 | 64,902 | 0 |
| | 2615840533B | 97.66 | 98.85 | 45,794 | 9.85 | 27.51 | 64,582 | 1 |
| | 2615840601 | 96.03 | 94.80 | 262,030 | 18.91 | 7.13 | 70,583 | 1 |
| | 2615840646 | 95.91 | 96.73 | 149,839 | 156.13 | 245.95 | 54,560 | 1 |
| | 2615840697 | 98.05 | 100.00 | 0 | 38.72 | 54.61 | 66,741 | 1 |
| | 2616644829 | 80.37 | 100.00 | 0 | 10.74 | 9.6 | 57,292 | 1 |
| | 2617270709 | 64.21 | 100.00 | 0 | 74.75 | 138.62 | 57,618 | 0 |
| | 2623620557 | 80.79 | 91.58 | 559,656 | 2430.04 | 837.43 | 12,423 | 1 |
| | 2623620567 | 86.45 | 94.83 | 354,741 | 1815.18 | 562.71 | 12,423 | 1 |
| | 2623620609 | 16.12 | 93.35 | 87,601 | 9557.15 | 902.32 | 12,423 | 0 |
| | 2623620617 | 79.31 | 64.06 | 1,471,566 | 1288.91 | 202.44 | 64,172 | 0 |
| | 2623620618 | 84.08 | 81.59 | 784,508 | 298.99 | 156.87 | 64,172 | 0 |
