## Supplemental Table 3 for "Hybrid Clustering of Long and Short-read for Improved Metagenome Assembly"

**Supplemental Table S3: Biosynthetic gene clusters predicted from alternative metagenome assembly protocols**

| Secondary metabolite class | Assemble-only | SpaRC-hybrid-Flye | SpaRC-hybrid-Canu |
|---|---|---|---|
| acyl_amino_acids | 5 | 1 | 1 |
| arylpolyene | 70 | 171 | 103 |
| bacteriocin | 259 | 480 | 244 |
| betalactone | 30 | 31 | 14 |
| butyrolactone | 1 | 0 | 1 |
| cyanobactin | 31 | 21 | 18 |
| ectoine | 13 | 9 | 0 |
| hglE-KS | 59 | 99 | 58 |
| hserlactone | 60 | 35 | 44 |
| indole | 46 | 78 | 37 |
| ladderane | 18 | 9 | 0 |
| lanthidin | 0 | 77 | 0 |
| lanthipeptide | 99 | 36 | 62 |
| LAP | 38 | 75 | 30 |
| lassopeptide | 98 | 73 | 41 |
| linaridin | 5 | 22 | 18 |
| microviridin | 5 | 26 | 10 |
| NAGGN | 13 | 5 | 11 |
| NRPS | 851 | 2259 | 1456 |
| nucleoside | 2 | 0 | 0 |
| oligosaccharide | 1 | 0 | 0 |
| other | 14 | 4 | 7 |
| PBDE | 6 | 0 | 0 |
| phenazine | 17 | 38 | 22 |
| phosphonate | 9 | 4 | 2 |
| PpyS-KS | 1 | 0 | 0 |
| proteusin | 6 | 6 | 0 |
| PUFA,hglE-KS | 0 | 1 | 2 |
| resorcinol | 36 | 98 | 56 |
| siderophore | 5 | 1 | 1 |
| T1PKS | 153 | 59 | 34 |
| T2PKS | 6 | 2 | 3 |
| T3PKS | 227 | 242 | 140 |
| terpene | 760 | 1398 | 729 |
| TfuA-related | 29 | 85 | 66 |
| thiopeptide | 12 | 7 | 5 |
| transAT-PKS | 3 | 6 | 2 |
| Total BGCs | 2988 | 5458 | 3217 |
